# The viral polymerase inhibitor 7-deaza-2’-*C*-methyladenosine is a potent inhibitor of *in vitro* Zika virus replication and delays disease progression in a robust mouse infection model

**DOI:** 10.1101/041905

**Authors:** Joanna Żmurko, Rafael E. Marques, Dominique Schols, Erik Verbeken, Suzanne J.F. Kaptein, Johan Neyts

**Author notes:** Address for correspondence: Johan Neyts, University of Leuven, Department of Microbiology and Immunology, Rega Institute for Medical Research, Laboratory of Virology and Chemotherapy, Minderbroedersstraat 10, B-3000 Leuven, Belgium.

## Abstract

Zika virus (ZIKV) is an emerging flavivirus typically causing a dengue-like febrile illness, but neurological complications, such as microcephaly in newborns, have potentially been linked to this viral infection. We established a panel of *in vitro* assays to allow the identification of ZIKV inhibitors and demonstrate that the viral polymerase inhibitor 7-deaza-2’-*C*-methyladenosine (7DMA) efficiently inhibits replication. Infection of AG129 (IFN-α/β and IFN-γ receptor knockout) mice with ZIKV resulted in acute neutrophilic encephalitis with viral antigens accumulating in neurons of the brain and spinal cord. Additionally, high levels of viral RNA were detected in the spleen, liver and kidney, and levels of IFN-γ and IL-18 were systematically increased in serum of ZIKV-infected mice. Interestingly, the virus was also detected in testicles of infected mice. In line with its *in vitro* anti-ZIKV activity, 7DMA reduced viremia and delayed virus-induced morbidity and mortality in infected mice, which also validates this small animal model to assess the *in vivo* efficacy of novel ZIKV inhibitors. Since AG129 mice can generate an antibody response, and have been used in dengue vaccine studies, the model can also be used to assess the efficacy of ZIKV vaccines.

**Article Summary Line:** A robust cell-based antiviral assay was developed that allows to screen for and validate novel inhibitors of Zika virus (ZIKV) replication. The viral polymerase inhibitor 7-deaza-2’-*C*-methyladenosine (7DMA) was identified as a potent ZIKV inhibitor. A mouse model for ZIKV infections, which was validated for antiviral studies, demonstrated that 7DMA markedly delays virus-induced disease in this model.

## Introduction

Zika virus (ZIKV), a mosquito-borne flavivirus, was first isolated from a febrile Rhesus monkey in the Zika Forest in Uganda in 1947 (1). During the last 5 decades sporadic ZIKV infections of humans were reported in Gabon, Nigeria, Senegal, Malaysia, Cambodia and Micronesia (2,3,4), leading to a benign febrile disease called Zika fever. The latter is characterized by headache, maculopapular rash, fever, arthralgia, malaise, retro-orbital pain and vomiting (5,6). In 2007, an epidemic of fever and rash associated with ZIKV infection was reported in Micronesia. During this outbreak 185 cases of ZIKV infections were confirmed. The seroprevalence in the affected region was 73% (7). During the more recent ZIKV outbreak in French Polynesia [FP] between October 2013 and February 2014 over 30,000 people sought medical care (8,9). Since then, ZIKV has spread to new areas in the Pacific, including New Caledonia, the Cook Islands, and Chile’s Easter Island (7,10). As of 2015 ZIKV is causing an epidemic in Central and South America with an increasing number of cases reported particularly in Brazil, Colombia and El Salvador (11-14), demonstrating that this is a truly emerging human pathogen. Hundreds of cases of Guillain-Barré syndrome have been reported in the wake of ZIKV infections (15,16,17). As a result of a marked increase in the number of cases of microcephaly among infants born to virus-infected women, Zika has been declared a public health emergency of national importance in Brazil (16,17,18). In addition, an increasing number of travelers returning sick from endemic regions were diagnosed with ZIKV (19-24). The *Aedes aegypti* mosquito, the primary vector for ZIKV transmission, is expanding in all (sub-)tropical regions of the world and was recently reported to be present in California, USA (25).

There is neither a vaccine nor a specific antiviral therapy for the prevention or treatment of infections by ZIKV. The increasing incidence of Zika fever stresses the need for both preventive and therapeutic measures. We here report on the establishment of (i) a panel of assays that allow to identify inhibitors of ZIKV replication as well as (ii) a robust animal model of ZIKV infection with brain involvement. The viral polymerase inhibitor 7-deaza-2’-*C*-methyladenosine (7DMA) was identified as an inhibitor of *in vitro* ZIKV replication and was shown to reduce viremia and to delay the time to disease progression in virus-infected mice.

## Materials and Methods

### Compounds

Ribavirin, 1-(β-D-ribofuranosyl)-1H-1,2,4-triazole-3-carboxamide (Virazole; RBV) was purchased from ICN Pharmaceuticals (Costa Mesa, CA, USA). 2’*-C-* methylcytidine (2’CMC) and 7-deaza-2’-*C*-methyl-D-adenosine (7DMA) were purchased from Carbosynth (Berkshire, UK). Favipiravir (6-fluoro-3-hydroxy-2-pyrazinecarboxamide; T-705) and its defluorinated analogue T-1105 (3-hydroxypyrazine-2-carboxamide) were obtained as custom synthesis products from abcr GmbH (Karlsruhe, Germany).

### Cells and viruses

ZIKV (strain MR766) was obtained from the European Virus Archive (EVA). Lyophilized virus was reconstituted in DMEM and virus stocks were generated on C6/36 mosquito cell cultures (ATCC^®^ CRL-1660™) grown in Leibowitz medium supplemented with 10% fetal calf serum (FCS), 1% non-essential amino acids (NEAA) and 20 nM HEPES at 28 °C, without CO2. At the time ZIKV caused a complete cytopathic effect (CPE) [d5-d7 post infection; pi] the supernatant was harvested and viral titers were determined by endpoint titration on Vero cells (African Green monkey kidney cells; ECACC), Vero E6 cells (Vero C1008; ATCC^®^ CRL-1586™) and BHK-J21 cells (baby hamster kidney cells; ATCC® CCL-10™). For end point titrations, cells were seeded in a 96-well plate at 5×10^3^ or 10^4^ cells/well in 100 μL assay medium and allowed to adhere overnight. The next day, 100 μL of ZIKV was added to each well, after which the virus was serially diluted (1:2). Following 5 days of incubation, culture medium was discarded and replaced with (3-(4,5-dimethylthiazol-2-yl)-5-(3-carboxymethoxyphenyl)-2-(4-sulfophenyl)-2H-tetrazolium; MTS) and the absorbance was measured at 498 nm following a 1.5h-incubation period. Subsequently, cultures were fixed with ethanol and stained with 1% Giemsa staining solution (solution of azure B/azure II-eosin/methylene blue 1:12:2 (w/w/w) in glycerol/methanol 5:24 (v/v); total dye content: 0.6 % (w/w) Sigma-Aldrich). The different cell types as well as ZIKV tested negative for mycoplasma.

### CPE-reduction assay

Vero cells were grown in growth medium, consisting of MEM (Life Technologies) supplemented with 10% FCS, 2 mM L-glutamine and 0.075% sodium bicarbonate (Life Technologies). Antiviral assays were performed using the same medium except that 10% FCS was replaced with 2% FCS, referred to as ‘assay medium’. Vero cells were seeded at a density of 10^4^ cells/well in a 96-well plate in 100 μL assay medium and allowed to adhere overnight. To each well 100 μl of culture medium containing 50% cell culture infectious doses (i.e., CCID_50_) of ZIKV was added, after which 2-fold serial dilutions of the compounds were added. Following 5 days of incubation CPE was determined by means of the MTS readout method and by microscopic evaluation of fixed and stained cells. In parallel, cell cultures were incubated in the presence of compound and absence of virus to evaluate a potential cytotoxic effect. The 50% effective concentration (EC_50_), which is defined as the compound concentration that is required to inhibit virus-induced CPE by 50%, and 50% cytotoxic concentration (CC_50_), which is defined as the compound concentration that is required to inhibit the cell growth by 50%, was visually determined. The Z’ factor was calculated by the following formula 1-[3×(SD_CC_+SD_VC_)/(OD_CC_-OD_VC_)]; VC, virus control; CC, cell control.

### Virus yield reduction assay

Vero cells were seeded at a density of 5×10^4^ cells/well in 96-well plates in growth medium and allowed to adhere overnight. Cells were washed 3 times with PBS and incubated with 100 μL CCID_50_ (MOI~0.2) of ZIKV in assay medium for 1 h at 37 °C. Next, cells were washed 3 times with PBS and 2-fold serial dilutions of the compounds were added. Supernatant was harvested at day 4 pi and stored at −80°C until further use. The EC_50_ value, which is defined as the compound concentration that is required to inhibit viral RNA replication by 50%, was determined using logarithmic interpolation.

### Viral kinetics and time-of-drug addition studies

Vero cells were seeded at a density of 2×10^5^ cells/well in 24-well plate in growth medium and allowed to adhere overnight. Cells were washed twice with PBS and incubated with ZIKV at an MOI~1 in assay medium for 30 min at 37 °C. After the incubation, cells were washed twice with PBS, after which assay medium was added to the cells. Cells were harvested at 0, 4, 6, 8, 10, 12, 14, 16, 18, 20, 22 and 24 hours pi and stored at −80°C until further use. For the time-of-drug addition studies, cells were seeded and infected as described above and 7DMA (178 μM) or ribavirin (209 μM) was added to the medium at different time points pi (see above). Cells were harvested at 24 hours pi and stored at −80°C until further use.

### Plaque reduction assay

Vero cells were cultured in growth medium. Cells were incubated with ZIKV for 1 h, washed and overlaid with a mixture of 2% (w/v) carboxymethylcellullose (Sigma Aldrich) and MEM supplemented with 2% FCS, 4 mM L-glutamine and 0.15% sodium bicarbonate. Two-fold serial dilutions of compounds were made in the overlay medium. Cells Cells were fixed and stained using a 10% v/v formaldehyde solution and a 1% methylene blue solution, respectively. Infectious virus titer (PFU/mL) was determined using the following formula: number of plaques × dilution factor × (1/inoculation volume).

### Immunofluorescence assay

Vero cells were infected with ZIKV as described for the virus yield reduction assay. After removal of the virus, 2-fold serial dilution (starting at 89 μM) of 7DMA was added to the cells. At 72 h pi, cells were subsequently fixed with 2% paraformaldehyde in PBS and washed with PBS supplemented with 2% BSA. Anti-Flavivirus Group Antigen Antibody clone D1-4G2-4-15 (Millipore) and goat anti-mouse Alexa Fluor 488 (Life Technologies) were used to detect ZIKV antigens in infected cells. Cell nuclei were stained using DAPI (4′,6-diamidino-2-fenylindool; Life Technologies) and read out was performed using an ArrayScan XTI High Content Analysis Reader (Thermo Scientific). The EC50 value, which is defined as the compound concentration that is required to inhibit viral antigen expression by 50%, was determined using logarithmic interpolation.

### RNA isolation and quantitative RT-PCR

RNA was isolated from 100-150 μl supernatant using the NucleoSpin RNA virus kit (Filter Service, Germany) according to the manufacturer’s protocol. RNA from infected cells was isolated using the RNeasy minikit (Qiagen, The Netherlands), according to the manufacturer’s protocol, and eluted in 50 μL RNase-free water. During RT-qPCR the ZIKV NS1 region (nucleotides 2472 - 2565) was amplified using primers 5’-TGA CTC CCC TCG TAG ACT G-3’ and 3’-CTC TCC TTC CAC TGA TTT CCA C-5’ and a Double-Quenched Probe 5’-6-FAM/AGA TCC CAC /ZEN/AAA TCC CCT CTT CCC/3’IABkFQ/ (Integrated DNA Technologies, IDT). Viral RNA was quantified using serial dilutions of a standard curve consisting of a synthesized gene block containing 145 bp of ZIKV NS1 (nucleotides 2456 - 2603): 5’-GGT ACA AGT ACC ATC CTG ACT CCC CTC GTA GAC TGG CAG CAG CCG TTA AGC AAG CTT GGG AAG AGG GGA TTT GTG GGA TCT CCT CTG TTT CTA GAA TGG AAA ACA TAA TGT GGA AAT CAG TGG AAG GAG AGC TCA ATG CAA TCC TAG-3’ (Integrated DNA Technologies).

### A mouse model of Zika virus infection

All experiments were performed with approval of and under the guidelines of the Ethical Committee of the University of Leuven [P087-2014]. Virus stock was produced as described earlier and additionally concentrated by ultracentrifugation. Infectious virus titers (PFU/ml) were determined by performing plaque assays on Vero cells. 129/Sv mice deficient in both interferon (IFN)-α/β and IFN-γ receptors (AG129 mice; male, 8-14 weeks of age) were inoculated intraperitoneally (ip; 200 μL) with different inoculums ranging from 1 ×10^1^ - 1× 10^5^ PFU/mL of ZIKV. Mice were observed daily for body weight change and the development of virus-induced disease. In case of a body weight loss of >20% and/or severe illness, mice were euthanized with pentobarbital (Nembutal). Blood was collected by cardiac puncture and tissues (spleen, kidney, liver and brain) were collected in 2-mL tubes containing 2.8 mm zirconium oxide beads (Precellys/Bertin Technologies) after transcardial perfusion using PBS. Subsequently, RLT lysis buffer (Qiagen) was added to the Precellys tubes and tissue homogenates were prepared using an automated homogenizer (Precellys24; Bertin Technologies). Homogenates were cleared by centrifugation and total RNA was extracted from the supernatant using the RNeasy minikit (Qiagen), according to the manufacturer’s protocol. For serum samples, the NucleoSpin RNA virus kit (Filter Service) was used to isolate viral RNA. Viral copy numbers were quantified by RT-qPCR, as described earlier.

## Histology

For histological examination, tissues (harvested at d13-15 pi) were subsequently fixed in 4% formaldehyde, embedded in paraffin, sectioned, and stained with hematoxylin-eosin, essentially as described before (26). Anti-Flavivirus Group Antigen Antibody, clone D1-4G2-4-15 (Millipore) was used to detect ZIKV antigens in tissue samples.

### Detection of pro-inflammatory cytokines and chemokines

Induction of pro-inflammatory cytokines and chemokines was analyzed in 20 μL serum using the mouse cytokine 20-plex antibody bead kit (ProcartaPlex Mouse Th1/Th2 & Chemokine Panel I [EPX200-26090-901]), which measures the expression of TNF-α, IFN-γ, IL-6, IL-18, CCL2 (MCP-1), CCL3 (MIP-1α), CCL4 (MIP-1β), CCL5 (RANTES), CCL7 (MCP-3), CCL11 (Eotaxin), CXCL1 (GRO-α), CXCL2 (MIP-2), CXCL10 (IP-10), GM-CSF, IL-1β, IL12p70, IL-13, IL-2, IL-4, and IL-5. Measurements were performed using a Luminex 100 instrument (Luminex Corp., Austin, TX, USA) and were analyzed using a standard curve for each molecule (ProcartaPlex). Statistical analysis was performed using a one-way ANOVA.

### Evaluation of the activity of 7DMA in ZIKV-infected AG129 mice

AG129 mice (male, 8-14 weeks of age) were treated with either 50 mg/kg/day 7DMA resuspended in 0.5% or 0.2% sodium carboxymethylcellulose (CMC-Na; n=9) or vehicle (0.5% or 0.2% CMC-Na; n=9) once daily (QD) via oral gavage for 10 consecutive days. Since the bulk-forming agent CMC has dehydrating properties (27), mice that received the drug (or vehicle) formulated with 0.5% CMC received (on days 6-9) subcutaneous injections with 200 μL of saline. One hour after the first treatment, mice were infected via the intraperitoneal route with 200 μL of a 1×10^4^ PFU/ml stock of ZIKV. Blood was withdrawn from the tails at different days pi. Viral RNA was extracted from 20 μL of serum using the RNA NucleoSpin RNA virus kit (Filter Service) followed by viral RNA quantification by means of RT-qPCR. Statistical analysis was performed using the Shapiro-Wilk normality test followed by the unpaired, two-tailed t-test in Graph Pad Prism6. Inter-group survival was compared using the Log-rank (Mantel-Cox) test. The *in vivo* efficacy of 7DMA was determined in two independent experimental animal studies. Evaluation of cytokine induction was performed using the ProcartaPlex Mouse Simplex IP-10 (CXCL10), TNF-α, IL-6 and IL-18 kits. In an additional animal study, AG129 mice (male, 8-14 weeks of age) were treated with 50 mg/kg/day 7DMA resuspended in 0.2% sodium carboxymethylcellulose (CMC-Na; n=6) or vehicle (0.2% CMC-Na; n=6) once daily (QD) via oral gavage for 5 successive days (starting 2 days prior to infection) and infected ip with 200 μL of a 1×10^4^ PFU/ml stock of ZIKV. Animals were euthanized at day 5 pi and testicles were collected and stored until further use.

## Results

### Establishing *in vitro* antiviral assays and the identification of 7DMA as a selective inhibitor of *in vitro* ZIKV replication

End point titrations in different cell lines revealed that Vero cells are highly permissive to ZIKV, hence, these cells were selected to establish antiviral assays. Infection with 100×TCID_50_ of ZIKV resulted in 100% cytopathic effect 5 days after infection (Supplementary Figure 1B), as assessed by microscopic evaluation as well as by the MTS readout method. The Z’ factor [a measure of statistical effect size to assess the quality of assays to be used for high-throughput screening purposes; (28)] of the CPE-reduction assay was 0.68 based on 64 samples (from 8 independent experiments) determined by the MTS readout method (Supplementary Figure 1C). The assay is thus sufficiently stringent and reproducible for high throughput screening purposes. The CPE-reduction assay was next employed to evaluate the potential anti-ZIKV activity of a selection of known (+)ssRNA virus inhibitors (i.e. 2’-*C*-methylcytidine, 7-deaza-2′-*C*-methyladenosine, ribavirin, T-705 and its analogue T-1105). All compounds resulted in a selective, dose-dependent inhibitory effect on ZIKV replication (Table 1). The antiviral effect of these compounds was confirmed in a virus yield reduction assay, a1.7log_10_ and 3.9log_10_ reduction in viral RNA load at a concentration of 22 μM and ≥45 μM, respectively, was noted (Table 1 and Figure 1A). Since 7DMA resulted in the largest therapeutic window (SI > 37; data not shown), the antiviral activity of this compound was therefore next assessed in a plaque reduction assay and in an immunofluorescence assay to detect viral antigens. The inhibitory effect of the compound in both assays was in line with those of the CPE-reduction and virus yield reduction assay (Table 1, Figure 1A). At a concentration of 11 μM, 7DMA almost completely blocked viral antigen expression (Figure 1B, left panel).

**Figure 1.**
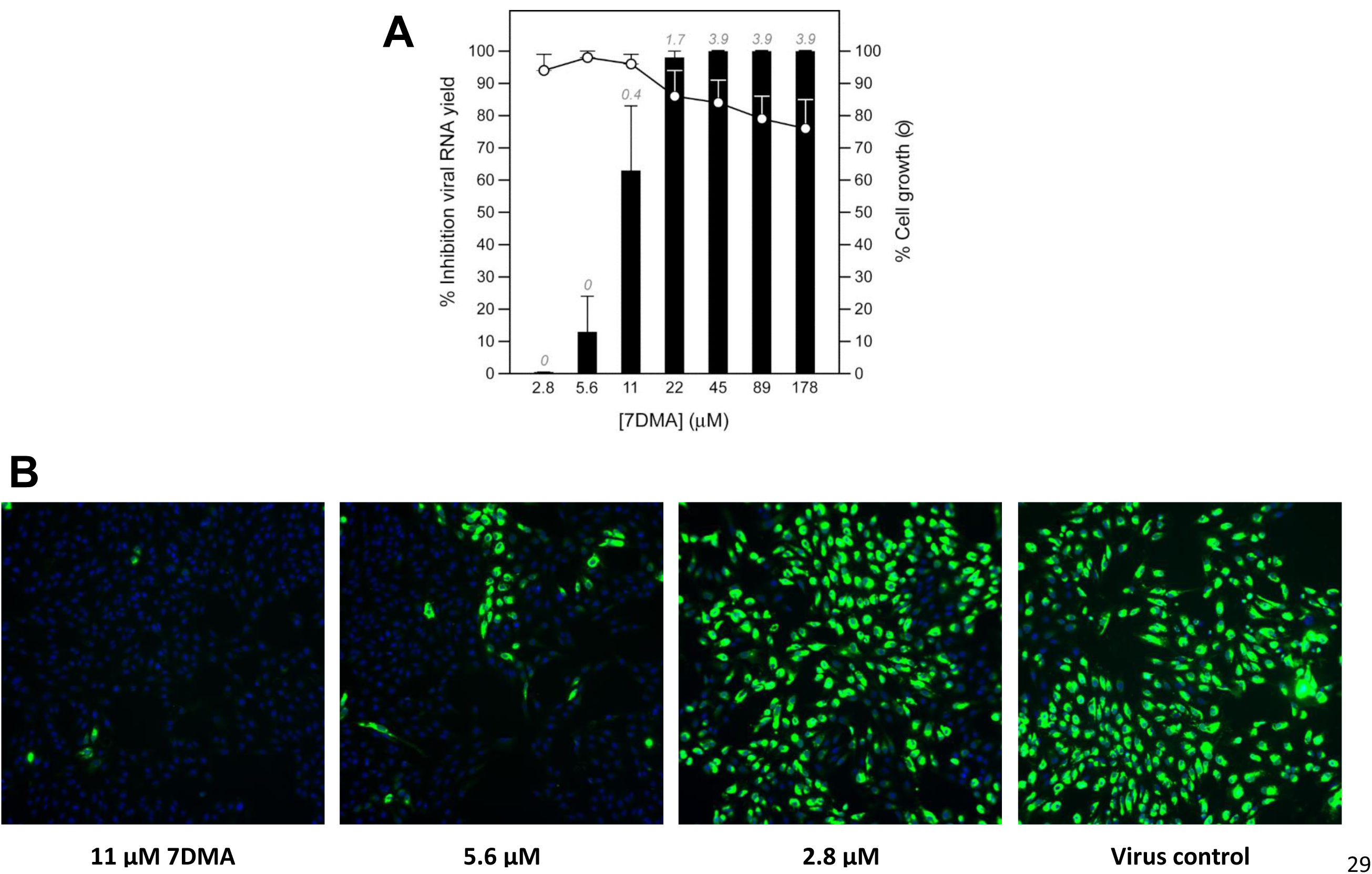
Dose-dependent inhibition of ZIKV RNA replication by 7DMA. (**A**) Vero cell cultures infected with ZIKV strain MR766 were treated with different concentrations of 7DMA. Viral RNA levels in the supernatant were quantified on day 4 pi by means of RT-qPCR and are expressed as percentage inhibition of untreated virus control (black bars). Mock-infected cells were treated with the same dilution series of 7DMA. Cell viability was determined by means of the MTS/PMS method and is expressed as percentage of cell growth of untreated control (white circles). Data represent mean values ± standard deviations (SD) for three independent experiments. Log10 reduction values in viral RNA load are depicted in italics at the top of each bar. (**B**) Antiviral activity of 7DMA against ZIKV as determined in an immunofluorescence assay. At a concentration of 11 μM, 7DMA almost completely blocked viral antigen expression (left panel) compared to untreated, infected cells (right panel) and infected cells treated at a lower concentration (5.6 and 2.8 μM; two panels in the middle).

**Table 1.** Antiviral and metabolic activity of a selection of compounds against ZIKV strain MR766 Antiviral activity was determined in a CPE reduction assay (CPE), virus yield reduction assay (VY), plaque reduction assay (PA), and immunofluorescence assay (IFA); metabolic activity was determined in a CPE reduction assay. Data represent median values ± standard deviations (SD) from two independent experiments with 2 replicates for each experiment (n=4), except for the result obtained in the PA. *NA*, not analyzed.

| Compound | EC<sub>50</sub> (μM) |  |  |  | CC<sub>50</sub> (μM) |
| --- | --- | --- | --- | --- | --- |
|  | CPE | VY | PA | IFA | CPE |
| Ribavirin | 13 ± 0.2 | 12 ± 6.6 | NA | NA | > 409 |
| T-705 | 22 ± 15 | 24 ± 7.0 | NA | NA | > 637 |
| T-1105 | 86 | 35 ± 15 | NA | NA | > 719 |
| 2'CMC | 10 ± 0.4 | 3.9 ± 1.6 | NA | NA | 28 ± 10 |
| 7DMA | 20 ± 15 | 9.6 ± 2.2 | 1.3 | 5.7 ± 2.2 | > 357 |

7DMA is, as its 5’-triphosphate metabolite, an inhibitor of viral RNA-dependent RNA polymerases. Addition of the compound to infected cells could be delayed until ~10 hours pi without much loss of antiviral potency; when first added at a later time point, the antiviral activity was gradually lost. This is line with the observation that onset of intracellular ZIKV RNA production was determined to occur at 10 to 12 hours pi (Figure 2). The reference compound ribavirin [a triazole nucleoside with multiple proposed modes of action; (29)], in contrast, already lost part of the antiviral activity when added at time points later than 4 hours pi (Figure 2).

**Figure 2.**
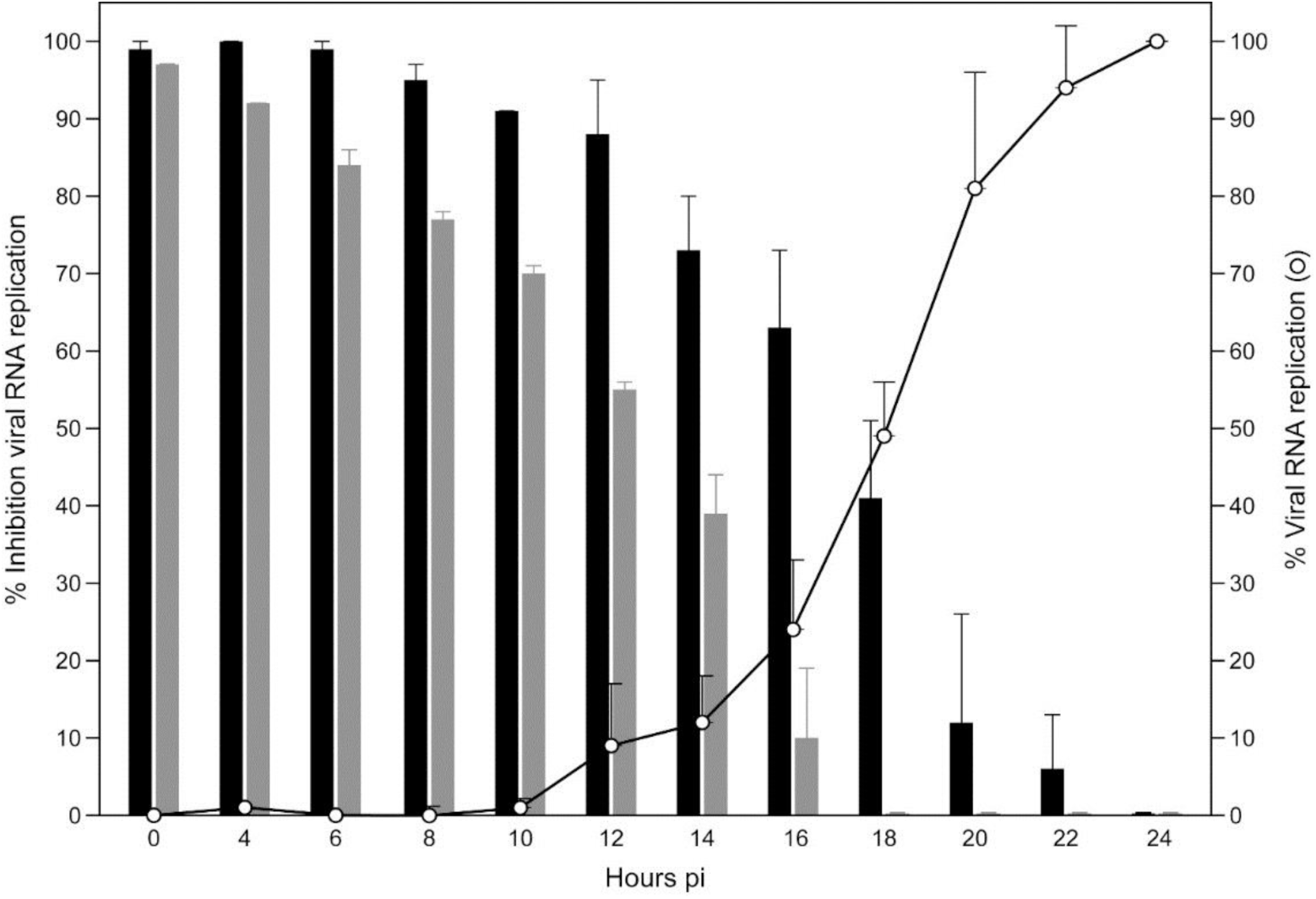
Viral replication kinetics of ZIKV and time-of-drug-addition studies. In viral kinetics studies, Vero cells were infected with ZIKV at an MOI~1.0 and harvested at the indicated time points pi. Data are expressed as percentage viral replication compared to viral RNA replication in infected cells at 24 hours pi (white circles). In time-of-drug-addition studies, ZIKV-infected cells were treated with 7DMA (178 μM; black bars) or ribavirin (205 μM; grey bars) at different time points pi. Cells were harvested at 24 hours pi and viral RNA was extracted and quantified by RT-qPCR. Data are expressed as percentage inhibition of viral replication compared to viral RNA replication in untreated, infected cells at 24 hours pi.

### Establishing a ZIKV infection model in mice

Intraperitoneal inoculation of IFN-α/β and IFN-γ receptor knockout mice (AG129) with as low as 200 μL of a 1 × 10^1^ PFU/ml stock of ZIKV resulted in virus-induced disease (see below) and mice had to be euthanized at a MDE (mean day of euthanasia) of 18.5 days pi (Figure 3A). Infection with higher inoculums (1 × 10^2^ - 1 × 10^5^ PFU/ml; 200 μL) resulted in a faster progression of the disease (MDE of 14 days pi) with the first signs of disease appearing at day 10 pi. Surprisingly, inoculation of SCID mice with 200 μL of a 1×10^4^ PFU/ml stock of ZIKV resulted in delayed disease; SCID mice had to be euthanized at day 40.0 ± 4.4 pi, roughly 26 days later than AG129 mice (data not shown). Disease signs in AG129 mice included paralysis of the lower limbs, body weight loss, hunched back and conjunctivitis. High levels of viral RNA were detected in brain, spleen, liver and kidney from mice that were euthanized at day 13-15 pi (Figure 3B). Histopathological analysis on tissues collected at day 13-15 pi revealed the accumulation of viral antigens in neurons of both the brain (Figure 4A) and the spinal cord (Figure 4D) as well as in hepatocytes (Figure 4E). Acute neutrophilic encephalitis (Figure 4C) was observed at the time of onset of virus-induced morbidity. It was next assessed whether infection with ZIKV resulted in the induction of a panel of 20 cytokines and chemokines at different time points pi (day 2, 3, 4 and 8; Figure 3C-3D and Supplementary Figure 2A-2G). In particular, levels of IFN-γ and IL-18 were increased systematically and significantly during the course of infection (Figure 3C and 3D), whereas levels of IL-6, CCL2, CCL5, CCL7, CXCL1, CXCL10 and TNF-α first increased, reaching a peak level at day 3 pi (CCL2, CXCL1, TNF-α Supplementary Figure 2A-2C) or day 4 pi (IL-6, CCL7, CXCL10; Supplementary Figure 2D-2F) pi and then gradually declined. Levels of CCL5 subsequently increased at day 2 pi, dropped at day 3 pi, and gradually increased again at day 4 and 8 pi (Supplementary Figure 2G).

**Figure 3.**
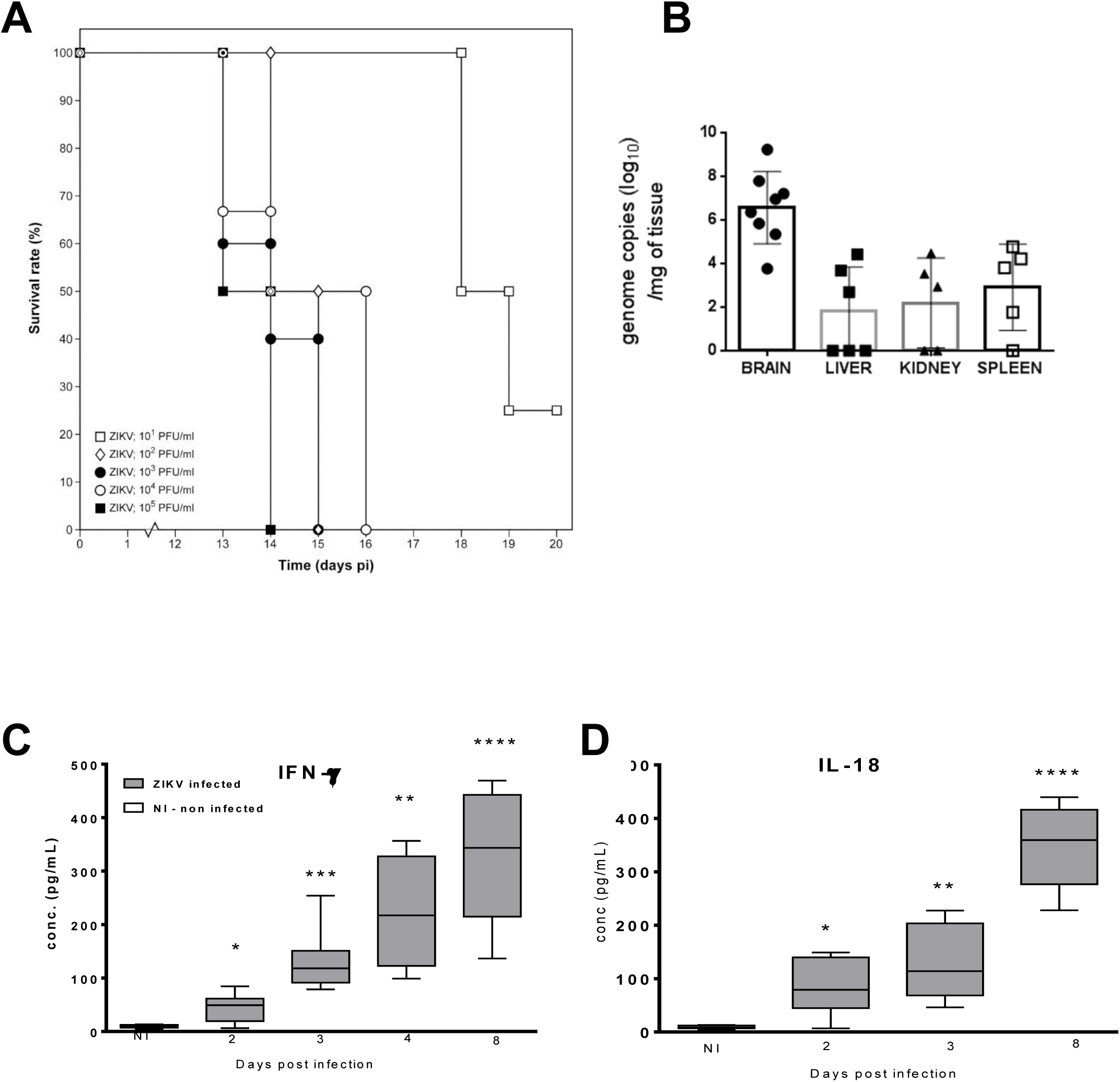
Establishment and characterization of an animal model for ZIKV infection. Male (8-14 weeks of age) 129/Sv mice deficient in both IFN-alpha/beta (IFN-α/β) and IFN-gamma (IFN-γ) receptors (AG129) were inoculated intraperitoneally with 200 μL of different inoculums (ranging from 1 × 10^1^ - 1× 10^5^ PFU/ml) of ZIKV. Mice were observed daily for body weight loss and the development of virus-induced disease. (**A**) Median day of euthanasia (MDE) is as follows: day 13.5, 15.0, 14.0, 14.5 and 18.5 pi for mice inoculated with 1×10^5^ (n=6), 1×10^4^ (n=6), 1×10^3^ (n=5), 1 ×10^2^ (n=2) and 1×10^1^ (n=4) PFU/mL, respectively. (**B**) Viral RNA load in brain (n=7), spleen (n=5), kidney (n=5) and liver (n=6) from ZIKV-infected mice as determined by RT-qPCR. Levels of IFN-γ (**C**) and IL-18 (**D**) were significantly increased throughout the course of infection in sera of AG129 mice (grey boxes) compared to those in sera of uninfected AG129 mice (white boxes). Statistical analysis was performed using the unpaired, two-tailed t-test. *, p<0.05.

**Figure 4.**
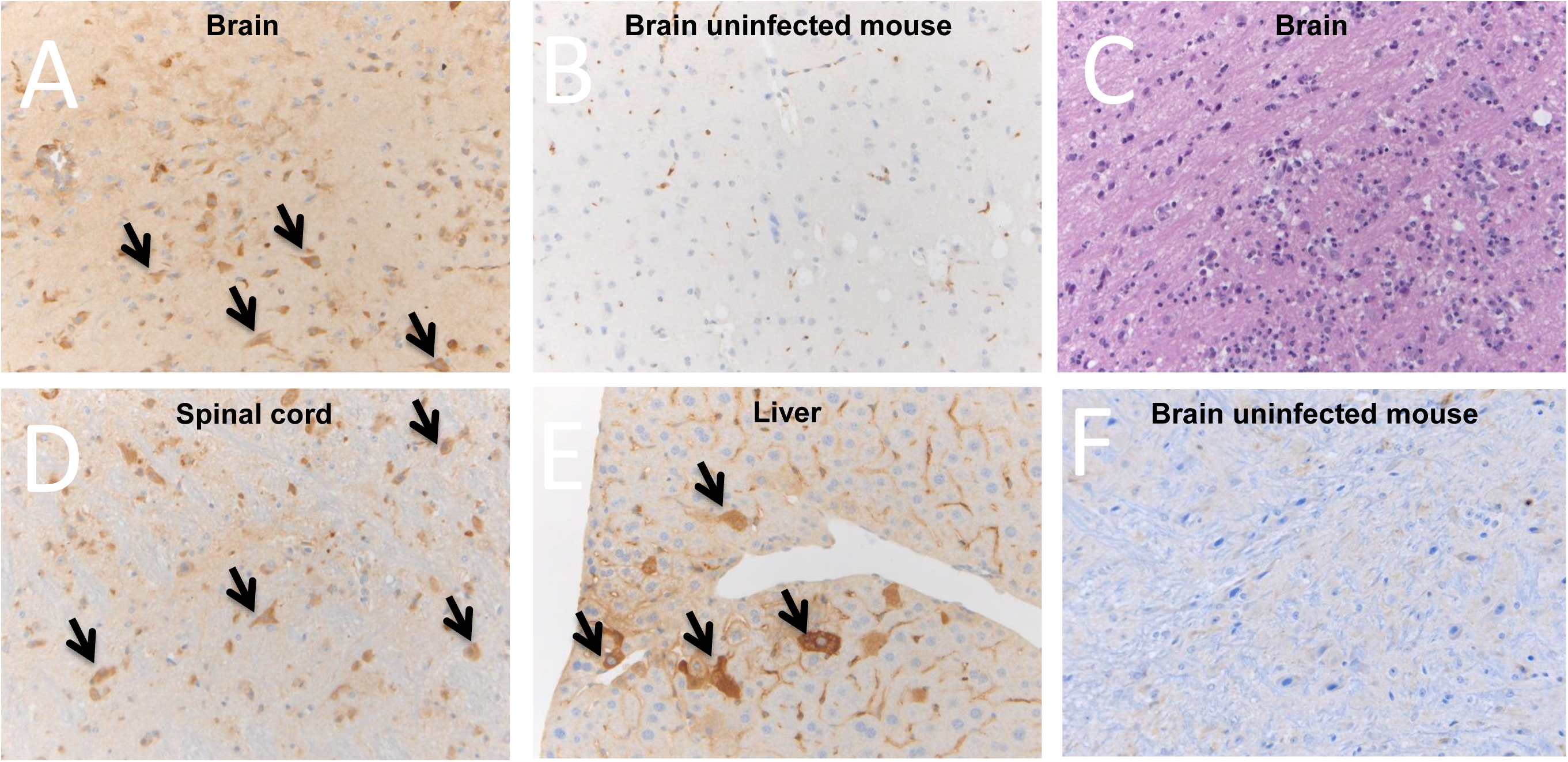
Presence of ZIKV antigens in the brain (**A**), spinal cord (**D**) and liver (**E**) of ZIKV-infected AG129 mice, whereas ZIKV antigens were absent in tissues of uninfected mice (brain, **B**), as shown by histopathological analysis. Infiltration of neutrophils is shown in the brain of ZIKV-infected mice (as detected by hematoxylin-eosin staining; **C**), but not in the brain of uninfected mice (**F**).

### 7-deaza-2’-*C*-methyladenosine delays ZIKV-induced disease in AG129 mice

AG129 mice were infected with 200 μL of a 1 × 10^4^ PFU/ml stock of ZIKV and were treated once daily with 50 mg/kg/day of 7DMA or vehicle *via* oral gavage (Figure 5) [data from the two independent experiments were not pooled since different amounts of CMC (respectively 0.5% and 0.2%) were used for formulation]. Vehicle-treated mice had to be euthanized two weeks after infection [MDE of 14.0 and 16.0 days, respectively]. 7DMA was well tolerated [no marked changes in body weight mass, fur, consistency of the stool or behavior during the treatment period] and markedly delayed virus-induced disease progression [MDE of 23.0 in the first study (p=0.003 as compared to the control) and 24.0 in the second study (p=0.04 as compared to the control)] (Figure 5A). 7DMA also reduced the viral RNA load in the serum of infected mice by 0.5log10, 0.8log10, 0.9log10, 0.7log10 and 1.3log10, respectively, at day 3, 5, 6, 7 and 8 pi (Figure 5B). Interestingly, at day 5 pi high levels of viral RNA (6.4log10) were found in the testicles of vehicle-treated mice (data not shown). At day 8 pi (shortly before the onset of disease in the vehicle controls), levels of IFN-γ in the serum were significantly higher in vehicle than in drug-treated mice (Figure 5C). No differences were noted in the expression of other cytokines between 7DMA-treated and untreated mice (data not shown).

**Figure 5.**
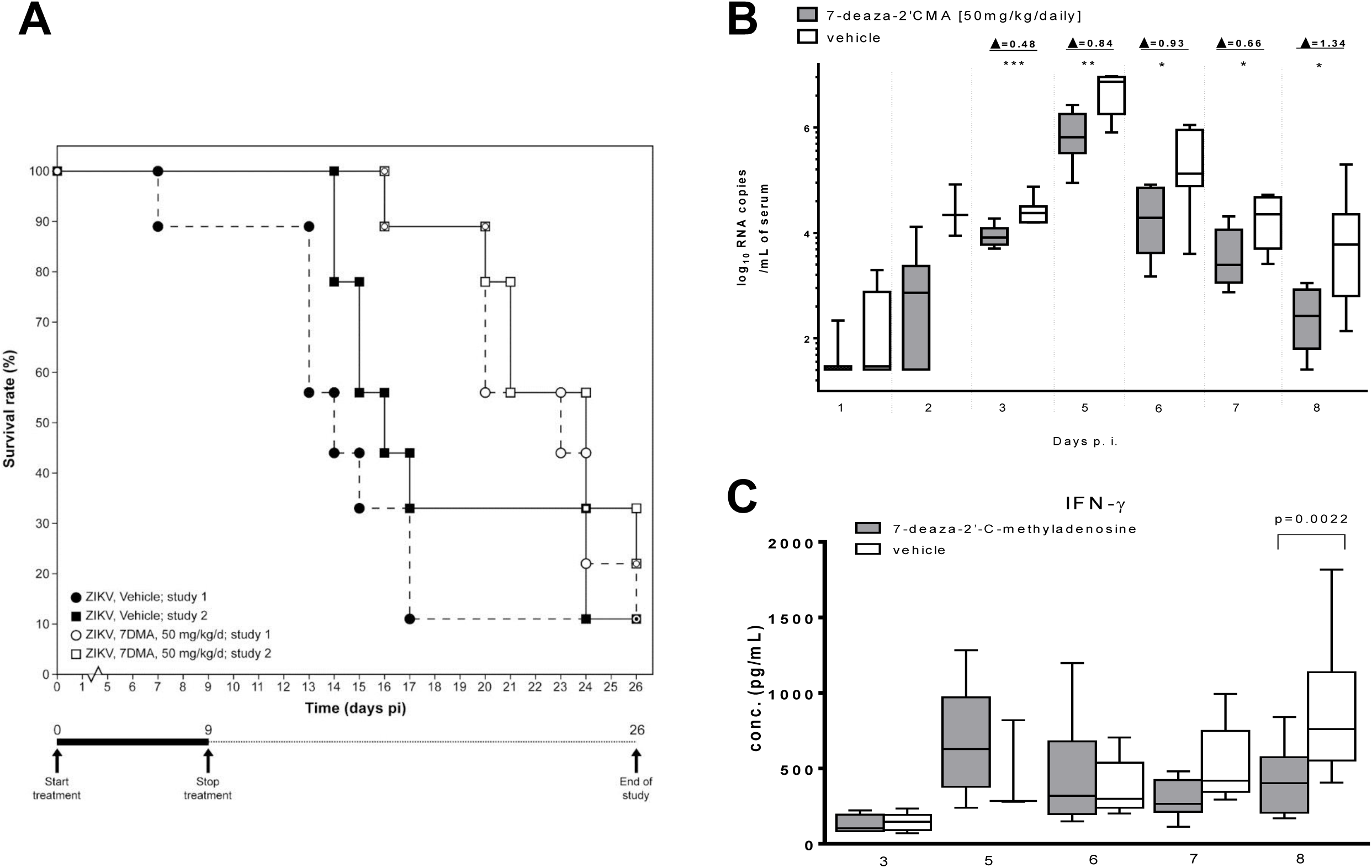
In vivo efficacy of 7DMA against ZIKV. AG129 mice (male, 8-14 weeks of age; n=9) were treated with 50 mg/kg/day 7DMA sodium carboxyinethylcellulose (CMC-Na)] via oral gavage or with vehicle [0.5% or 0.2% CMC-Na; n=9] for 10 days. Mice were infected intraperitoneally with 200 μL of a 1 × 104 PFU/inL stock of ZIKV 1 hour after the first treatment on day 0. (**A**) Percentage survival between ZDCV-infected mice treated with vehicle (• and ■) or 7DMA (○ and □) was compared using the Log-rank (Mantel-Cox) test. Data represent results from 2 independently performed studies. (**B**) Viral RNA load in seruni on day 1, 2, 3, 5, 6, 7 and 8 pi of ZDCV-infected mice treated with vehicle (white boxes) or 7DMA (grey boxes), as deternuned by RT-qPCR. Statistical analysis was performed using the unpaired, two-tailed t-test. Data are representative of 2 independent experiments. (**C**) Expression at different time points pi of IFN-γ in sera of ZIKV-infected mice treated with vehicle (white boxes) or 7DMA (grey boxes), as determined using the ProcartaPlex Mouse IFN-γ, IL-18, IL-6, IP-10, TNF-α Simplex kit (e-Bioscience). Data represent results from 2 independent experiments.

## Discussion

The rapid geographical spread of ZIKV, particularly in Central and South America poses a serious public health concern given that infection with this virus is less benign than initially thought. Hundreds of patients have been reported with Guillain-Barré syndrome (16,17). Most importantly, in Brazil a dramatic upsurge in the number of cases of microcephaly has been noted in children born to mothers infected with ZIKV. The annual rate of microcephaly in Brazil has increased from 5.7 per 100 000 live births in 2014 to 99.7 per 100 000 in 2015 (16,17,18). There is, hence, an urgent need to develop preventive and counteractive measures against this truly neglected flavivirus member. We here report on the establishment of (i) *in vitro* assays that will allow to identify novel inhibitors of ZIKV replication and (ii) a ZIKV infection model in mice in which the potential efficacy of such inhibitors can be assessed. ZIKV was found to replicate efficiently in Vero cells and to produce full CPE within a couple of days. The Z’ factor that was calculated for a colorometric (MTS method) CPE-based screen indicated that this is a robust assay that is amenable for high-throughput screening purposes. A plaque reduction, an infectious virus yield and a viral RNA yield reduction assay as well as an immunofluorescent antigen detection assay were established that will allow to validate the *in vitro* activity of hits identified in CPE-based screenings. Productive infection of human dermal fibroblasts, epidermal keratinocytes and immature dendritic cells with the ZIKV has recently been reported (30). However, Vero cells may be ideally suited for high throughput screening purposes, making these cells most useful to confirm the antiviral activity of interesting inhibitors of viral replication. We employed the assays that we established to assess the potential anti-ZIKV activity of a number of molecules with reported antiviral activity against other ssRNA viruses. In particular, the nucleoside analogue 7DMA was identified to inhibit ZIKV replication with a potency that was more or less comparable between the different *in vitro* assays. 7DMA was originally developed by Merck Research Laboratories as an inhibitor of hepatitis C virus replication (31), but was also shown to inhibit the replication of multiple flaviviruses, [i.e. dengue virus, yellow fever virus as well as West Nile and tick-borne encephalitis virus] with EC50 values ranging between 5 and 15 μM, which is thus comparable to the EC_50_ values for inhibition of *in vitro* ZIKV replication (31,32,33). In line with its presumed mechanism of action, i.e. inhibition as its 5’-triphosphate of the viral RNA-dependent RNA polymerase, time-of-drug-addition experiments revealed that the compound acts at a time point that coincides with the onset of intracellular viral RNA replication.

To assess the *in vivo* efficacy of ZIKV inhibitors, we established a model of ZIKV infection in mice. AG129 mice proved highly susceptible to ZIKV infections; even an inoculum of ~10 PFU/ml resulted in virus induced-morbidity and mortality. Although ZIKV-infected SCID mice (deficient in both T and B lymphocytes) developed severe disease requiring euthanasia (data not shown), these mice were more resistant to ZIKV infection than AG129 mice. SCID mice succumbed to infection roughly 26 days later than AG129 mice when inoculated with the same viral inoculum. Thus, ZIKV infections in mice are mostly by the interferon response rather than by lymphocytes, indicating that the innate immune response to ZIKV is critical. AG129 mice have been shown to be highly susceptible to infection with other flaviviruses; in particular allowing the development of dengue virus infection models in mice (32,34,35). At the time of virus-induced morbidity and mortality, ZIKV was detected in multiple organs such as kidney, liver and spleen, but also in the brain and spinal cord. The latter is in line with the observation that infected mice developed acute neutrophilic encephalitis with movement impairment and paralysis of the limbs. Brain involvement in ZIKV-infected mice may be relevant for brain-related pathologies in some ZIKV-infected humans (16,17). Interestingly, the virus was also detected at high levels in the testicles of infected mice. A few cases of sexual transmission of the ZIKV in humans have been reported (36,37); the observation that the virus replicates in the testicles in mice may suggest that the virus can also replicate in human testicle tissue thus explaining sexual transmission.

Pro-inflammatory cytokines (IFN-γ, IL-18, IL-6, TNF-α) and chemokines (CCL2, CCL5, CCL7, CXCL1, CXCL10) were found to be increased in sera of ZIKV-infected mice, indicating that infection causes systemic inflammation. In particular IFN-γ and IL-18 were continuously increased during the course of infection; both cytokines could therefore potentially function as predictive markers of disease progression and disease severity in this mouse model. Whether these cytokines are also upregulated during the acute phase of the infection in humans remains to be studied. Of note, the fact that ZIKV infection leads to the production of IL-18 suggests that the inflammasome is activated during the course of infection. Surprisingly, we could detect increased levels of IL-18, but not of IL-1β, which is also produced upon activation of the inflammasome (38). To our knowledge, the observation that the inflammasome could be implicated in ZIKV infection is unprecedented.

Recently, a small study was reported involving 6 ZIKV-infected patients in which during the acute phase 11 cytokines/chemokines were found to be significantly increased, of which 7 were also increased during recovery (39). Despite the fact that immunocompromised AG129 mice have an altered cytokine metabolism and were infected with the prototype ZIKV MR766 strain belonging to a different lineage than the one infecting the Latin American patients (African versus Asian, respectively), similarities in cytokine expression were noted between both studies. IL-6, CCL5 and CXCL10 were significantly increased in ZIKV-infected patients as well as in the infected mice. In the ZIKV-infected patients IFN-γ levels, which were markedly increased in ZIKV-infected mice, were also increased during both the acute and the reconvalescent phase of the infection, albeit non-significantly. Likewise, TNF-α levels, which were increased early in infection in mice, were (non-significantly) increased during the acute phase of infection in the patients. More studies are necessary to assess whether the cytokine profile in these 6 patients is representative for larger groups.

Treatment of ZIKV-infected mice with 7DMA significantly reduced viremia (between day 3 and 8 post infection) and delayed virus-induced morbidity and mortality. The compound was very well tolerated in mice, which is in line with earlier reports (31). The reduction in viremia and, hence, the delay of virus-induced disease was relatively modest, which is not surprising given the relatively weak *in vitro* activity of the compound as compared to, for example, the EC_50_ values (sub μM or even nM range) of most HCV inhibitors. Most importantly, the use of this compound allowed to validate the ZIKV mouse model to assess the efficacy of ZIKV inhibitors. Whether 7DMA (or related analogues) may have future in the control of ZIKV infections remains to be explored. AG129 mice have been used as well in the development of DENV vaccines, the DENV AG120 mouse models offer multiple disease parameters to evaluate protection by candidate vaccines (40). Hence, the ZIKV mouse model presented here may also serve to study the efficacy of vaccine strategies against the ZIKV.

In conclusion, we here report on a panel of *in vitro* cellular assays that will allow to run large-scale antiviral screening campaigns against ZIKV and to validate the antiviral activity of hit compounds. A number of molecules, including the viral polymerase inhibitor 7DMA, were found to inhibit the *in vitro* replication of ZIKV. Hence, 7DMA can be used as a reference compound/comparator in future studies. Moreover, a robust ZIKV mouse infection model was established; 7DMA delayed virus-induced mortality and, hence, validates this model for antiviral studies. Moreover, the model may be useful to study the efficacy of vaccination strategies against the ZIKV.

## Acknowledgements

This work was funded by EU FP7 [FP7/2007-2013] project EUVIRNA under grant agreement n° [264286], by EU FP7 SILVER (contract no. HEALTH-F3-2010-260644), a grant from the Belgian Interuniversity Attraction Poles (IAP) phase VII – P7/45 (BELVIR), and the Fondation Dormeur, Vaduz. Part of this research work was performed using the ‘Caps-It’ research infrastructure (project ZW13-02) that was financially supported by the Hercules Foundation and Rega Foundation, KU Leuven. We thank Els Vanstreels, Carolien De Keyzer, Natasha Danda, Sandra Claes, Mareike Grabner and Ruben Pholien for excellent technical assistance.

---

**Supplementary Figure 1.**
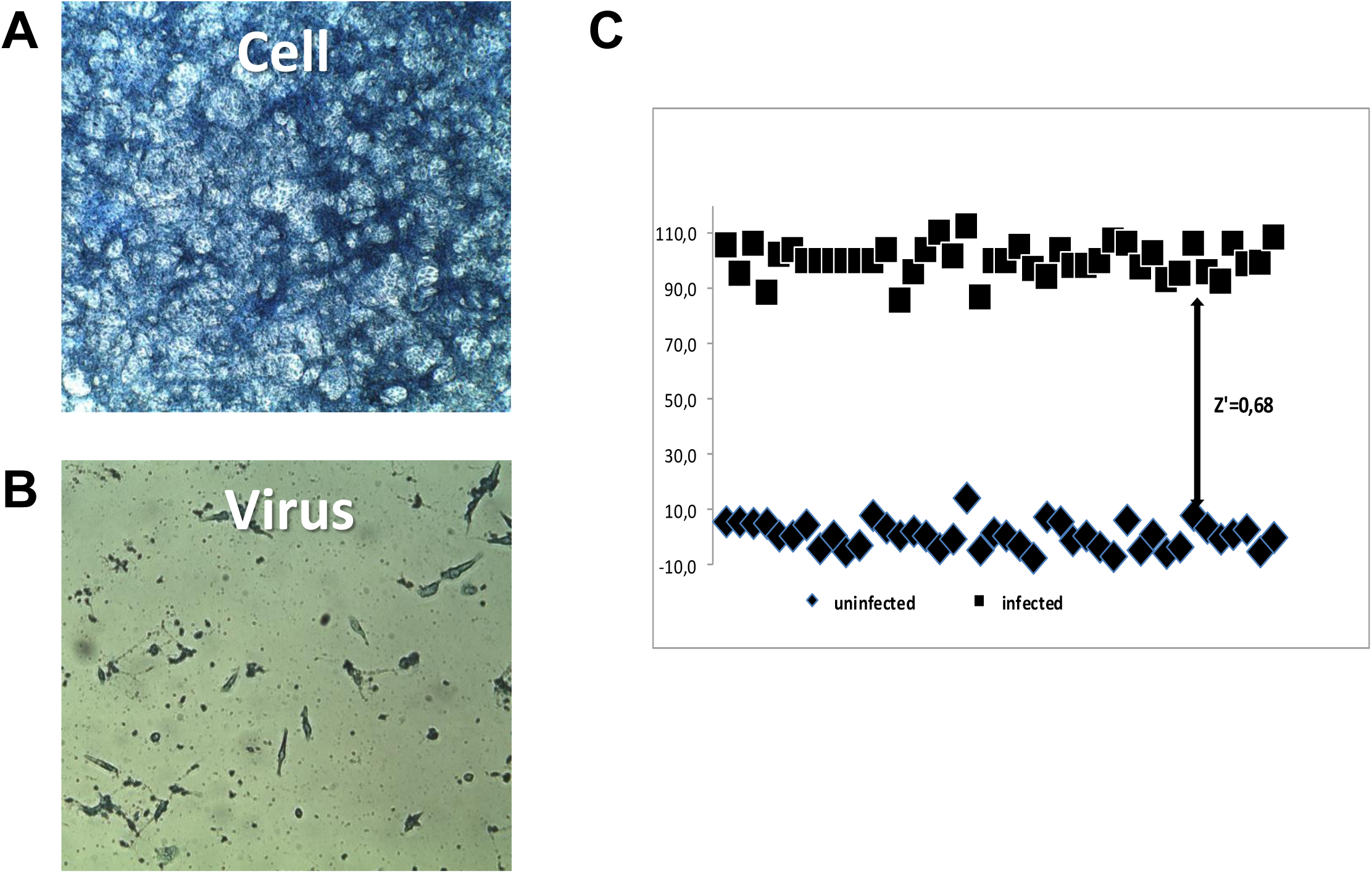
CPE reduction assay. Vero cells infected with ZIKV MR766 causes full CPE (**B**) at day 5 pi; uninfected cells (**A**). Z’ factor (0.68) was calculated for 64 samples (in 8 independent experiments; **C**) determined by the MTS readout method using the formula: 1-[3×(SD_CC_+SD_VC_)/(OD_CC_-OD_VC_)]; VC, virus control; CC, cell control.

**Supplementary Figure 2.**
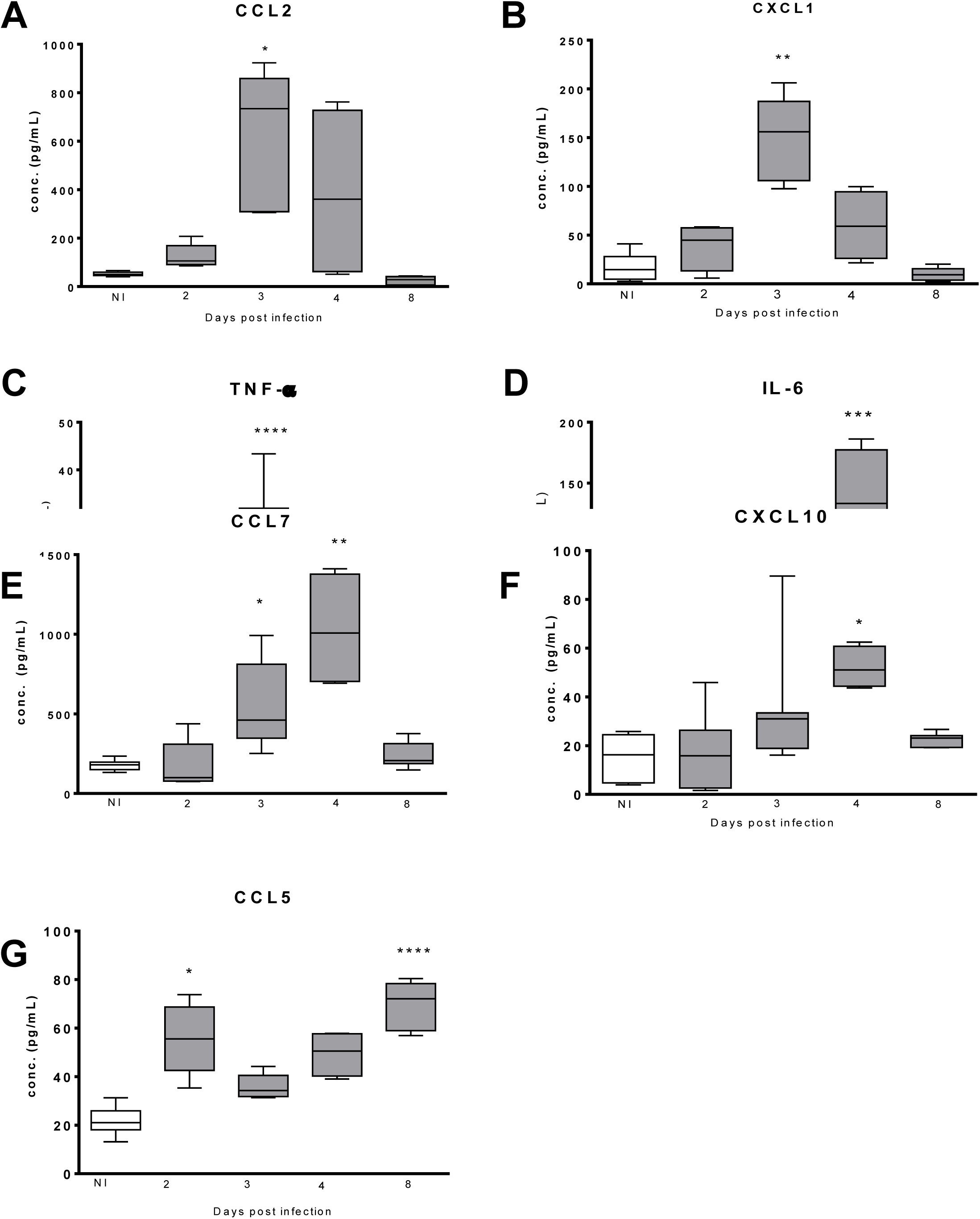
Box-and-whiskers plots showing increased levels of a panel of cytokines and chemokines that were increased in sera of ZIKV-infected mice (grey boxes) at different days pi compared to those in sera of non-infected mice (white boxes): (**A**) CCL2, (**B**) CXCL1, (**C**) TNF-α, (**D**) IL-6, (**E**) CXCL10, **F**) CCL5 and (**G**) CCL7. Induction of cytokines and chemokines was detected in 20 μL of serum using the ProcartaPlex™ Multiplex Immunoassay Panel with Mouse Th1/Th2 & Chemokine Panel 20-Plex (e-Bioscience). Statistical analysis was performed using a one-way. *, p<0.05.

